# Impact of PI3K pathway alterations on response to immune checkpoint inhibitors in HPV-negative head and neck squamous cell carcinoma

**DOI:** 10.1101/2024.07.02.601738

**Authors:** Jong Chul Park, Nandini Pal Basak, Mateo Useche, Jun Seok Park, Mandakulutur Subramanya Ganesh, Amritha Prabha, Chandra Reddy JayaPrakash, Mark Varvares, Lori Wirth, William Faquin, Satish Sankaran, Srinivas Vinod Saladi

## Abstract

Phosphatidylinositol-3-kinase (PI3K)/AKT/mammalian target of rapamycin (mTOR) pathway (PI3K pathway) is a major intracellular regulatory pathway commonly involved in cancer survival, proliferation, migration, and metabolism. Activating mutations and amplifications of phosphatidylinositol-4,5-bisphosphate 3-kinase catalytic subunit alpha (*PIK3CA*) are frequent genomic features of head and neck squamous cell carcinoma (HNSCC). A growing body of evidence suggests that dysregulation of the PI3K pathway suppresses the anti-tumor immune response, thereby enabling tumor immune evasion. We retrospectively analyzed clinicopathologic and genomic data of patients with recurrent or metastatic (R/M) HNSCC treated with immune checkpoint inhibitors (ICI). PI3K pathway alterations were found in 44% and were associated with poor clinical outcomes in the human papillomavirus-negative (HPV-ve) HNSCC patients. *PIK3CA* expression was found to be inversely correlated with immune gene expression, decreased T-cell infiltration, and reduced CD8 T-cell cytotoxic activity. The pharmacologic inhibition of the PI3K pathway induced an increase in key immune-responsive gene expression in *PIK3CA-*mutant HPV-ve HNSCC cells. These findings suggest that the PI3K pathway activation promotes immune suppressive phenotype, and playing a role in conferring resistance to immune checkpoint inhibitors in HPV-ve subset of HNSCC patients.

**Highlights:**

Alterations in PI3K pathway are associated with poor clinical outcomes in HPV-negative HNSCC patients treated with immune checkpoint therapy.
PI3K pathway activation correlates to immune suppressive tumor microenvironment in HNSCC.
Pharmacologic inhibition of PI3K induces the expression of interferon-gamma responsive and antigen presentation machinery genes in HNSCC.

## INTRODUCTION

Head and neck squamous cell carcinoma (HNSCC) is the 6^th^ most common type of cancer worldwide, with 900,000 annual new cases^1,2^. About 60% of patients with HNSCC present with locally advanced disease, and more than half of patients develop recurrence or metastasis (R/M)^3^. Despite advances in therapeutics, the prognosis of R/M HNSCC remains poor, with a median survival of around 12 months^4^. While immunotherapy with immune checkpoint inhibitors (ICI) targeting programmed cell death protein-1 (PD-1) has demonstrated highly effective and durable clinical benefit for patients with R/M HNSCC^5^, only a small portion of patients benefit from ICI with a response rate of 15-20%^4,6,7^. Suboptimal response to ICI in HNSCC may be attributed to various factors, including tumor intrinsic genomic, epigenetic, and metabolic features, and hostile tumor immune microenvironment created by immune suppressive immune cells, tumor-promoting cytokines, and dysregulated tumor stroma and vasculature^5,8^. Identifying predictive biomarkers may enable optimal patient selection and help to develop rational combinational therapeutic treatment strategies to overcome resistance ^9,10^.

The phosphatidylinositol-3-kinase (PI3K)/AKT/mammalian target of rapamycin (mTOR) pathway (PI3K pathway) is among the most dysregulated intracellular pathways in solid cancers^11–13^. Within this context, Class IA PI3K catalyzes the conversion of phosphatidylinositol 4,5-biphosphate (PIP2) to phosphatidylinositol (3,4,5)-triphosphate (PIP3) at the cellular membranes upon the binding of growth factors to receptor tyrosine kinases^14^. These lipid molecules act as second messengers, triggering a signaling cascade that regulates cellular functions, including survival, cell growth, proliferation, migration, and metabolism. The phosphatidylinositol-4,5-bisphosphate 3-kinase catalytic subunit alpha (*PIK3CA*) gene encodes the catalytic subunit phosphoinositide 3-kinase alpha protein, a critical component of the PI3K pathway. PI3K pathway alterations, including activating mutations and amplifications, are found in over 30% of HNSCC and are associated with poor prognosis^15–18^. A growing body of evidence including studies from mouse models suggests PI3K pathway dysregulation is implicated in antagonizing anti-tumor immune responses, thereby enabling tumor immune evasion^19–22^. However, the underlying mechanisms behind these effects have not been elucidated, and their effects on immunotherapy response in patients remain largely unexplored in the context of HPV-ve HNSCC patients^19,23^. In the present study, we utilize genomic analysis of patients who received ICI and report novel findings suggesting that PI3K pathway alterations may adversely affect the response HPV-ve HNSCC to ICI.

## RESULTS

### Clinicopathologic and genomic characteristics

A retrospective analysis was conducted on clinical data from patients with R/M HNSCC treated with anti-PD1-based ICI at Massachusetts General Cancer Center who had available next-generation sequencing (NGS)-based molecular test information. The analysis included 93 patients who met the selection criteria (see Methods section for details). The median age was 65, with 77% being male. Of the total cases, 33% were human papillomavirus (HPV)-positive (+ve), and 67% were HPV-negative (-ve), or unknown. All unknown cases were of non-oropharyngeal origin. All patients received anti-PD1 ICI, either pembrolizumab or nivolumab. Fifty-five (59%) patients received ICI as 1^st^-line (1L) therapy, and most patients (80%) had ICI as a monotherapy. Nineteen patients received a combination ICI with an anti-PD1 antibody and experimental immunotherapy agent through clinical trials (ClinicalTrail.gov Identifier: NCT04675294, NCT03978689, NCT03809624, NCT04398524, NCT02488759). The median PD-L1 combined positive score (CPS) was 10. Further baseline demographic and clinicopathologic information are summarized in Table S1.

The NGS-based sequencing analysis revealed mutations in a total of 75 genes in this cohort of patients (Figure S1). *TP53* (43%), *CDKN2A* (27%), *PIK3CA* (27%), *TERT* (24.7%), and *NOTCH1* (15%) were the most frequently altered genes (Table S2). Notably, *PIK3CA* was the most frequently altered gene (35%) (Table S2) in HPV+ve patients. Analysis of The Cancer Genome Atlas (TCGA) also showed amplification of PIK3CA gene along with missense mutations which are putative drivers (Figure S2A). A signaling pathway-centered analysis enriched for alterations in genes related to cell cycle control (52.7%), PI3K signaling pathway (44.1%), MAP kinase signaling (36.6%), and cell differentiation (30%) as the most prevalent (Figure 1 and Table S3).

**Figure 1.**
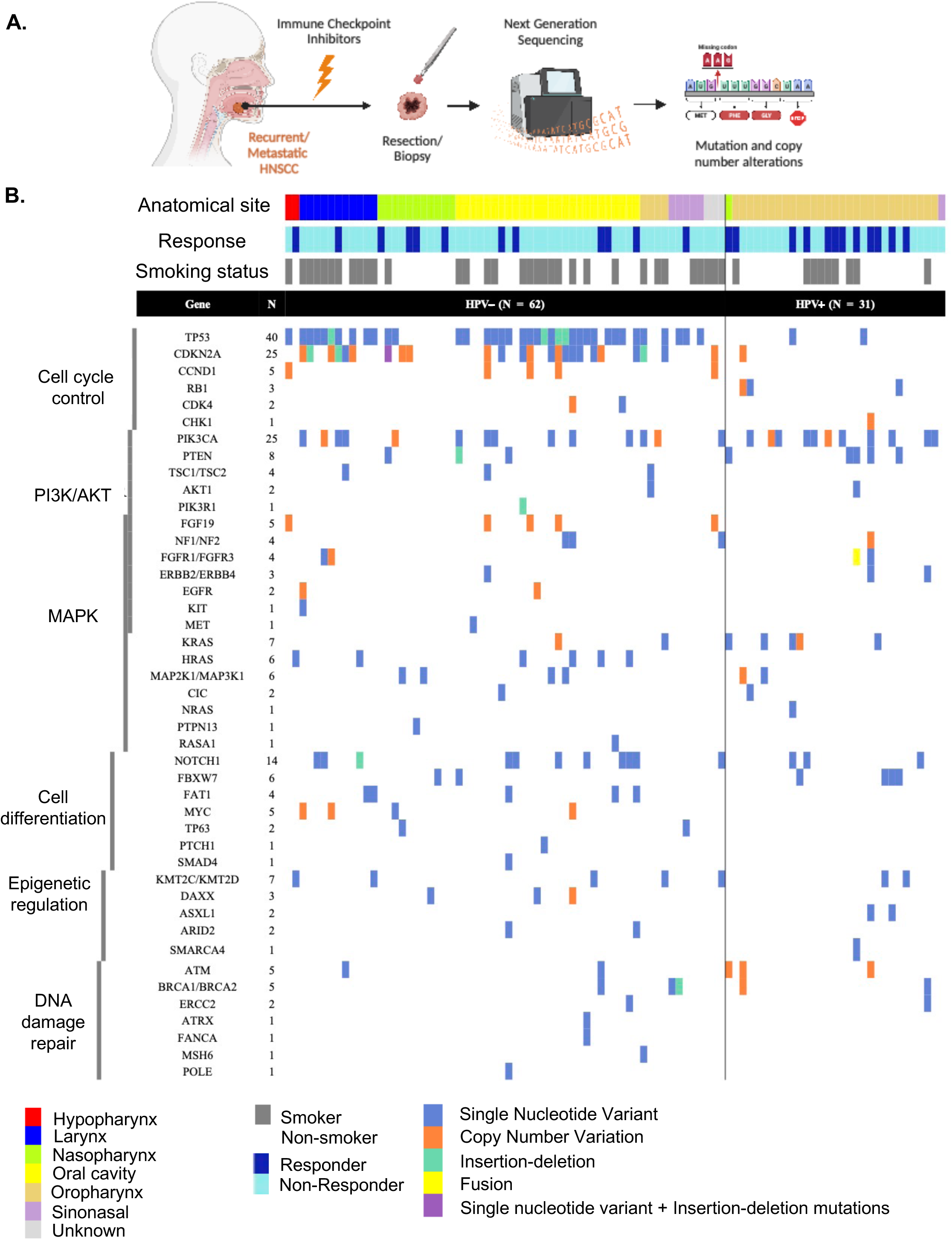

With a median follow-up of 14.2 months, the median overall survival (OS) and progression-free survival (PFS) were 18.3 and 3.8 months, respectively. Radiographic response (defined as the unequivocal decrease of overall tumor size(s) regardless of the magnitude) was seen in 26% of patients. Clinicopathologic factors which correlated with survival include a radiographic response (responder vs. non-responder) (HR 0.17, *p* < 0.001) and ECOG status (1/2 vs. 3/4) (HR 0.28, P < 0.001). Survival trends related to PD-L1 (CPS ≥ 10 vs. < 10, *p* = 0.82) did not meet statistical significance.

### Association between PI3K pathway alteration and clinical outcomes

Patients with PI3K pathway alterations exhibited a shorter survival trend, and this inferior survival was statistically significant in the HPV-ve population (20.9 vs. 13.0 months, HR 2.01, 95% CI 1.01 – 3.98, *p* = 0.05) (Figure 2A and 2B). Similarly, the PI3K pathway alteration was associated with an inferior PFS trend in the HPV-ve population (4.8 vs. 3.4 months, HR 1.87, *p* = 0.04) (Figure 2A and 2B). Patients with altered PI3K pathway also had a lower radiographic response rate compared to the PI3K pathway non-altered group in the HPV-ve population, 3.8% vs. 30.6%, respectively. In summary, our findings suggest a negative association between PI3K pathway alteration with survival outcomes as well as response to ICI in the HPV-ve HNSCC population, indicating a potential role of the PI3K pathway in mediating resistance to immunotherapy and poorer clinical benefit in HPV-ve HNSCC patients.

**Figure 2.**
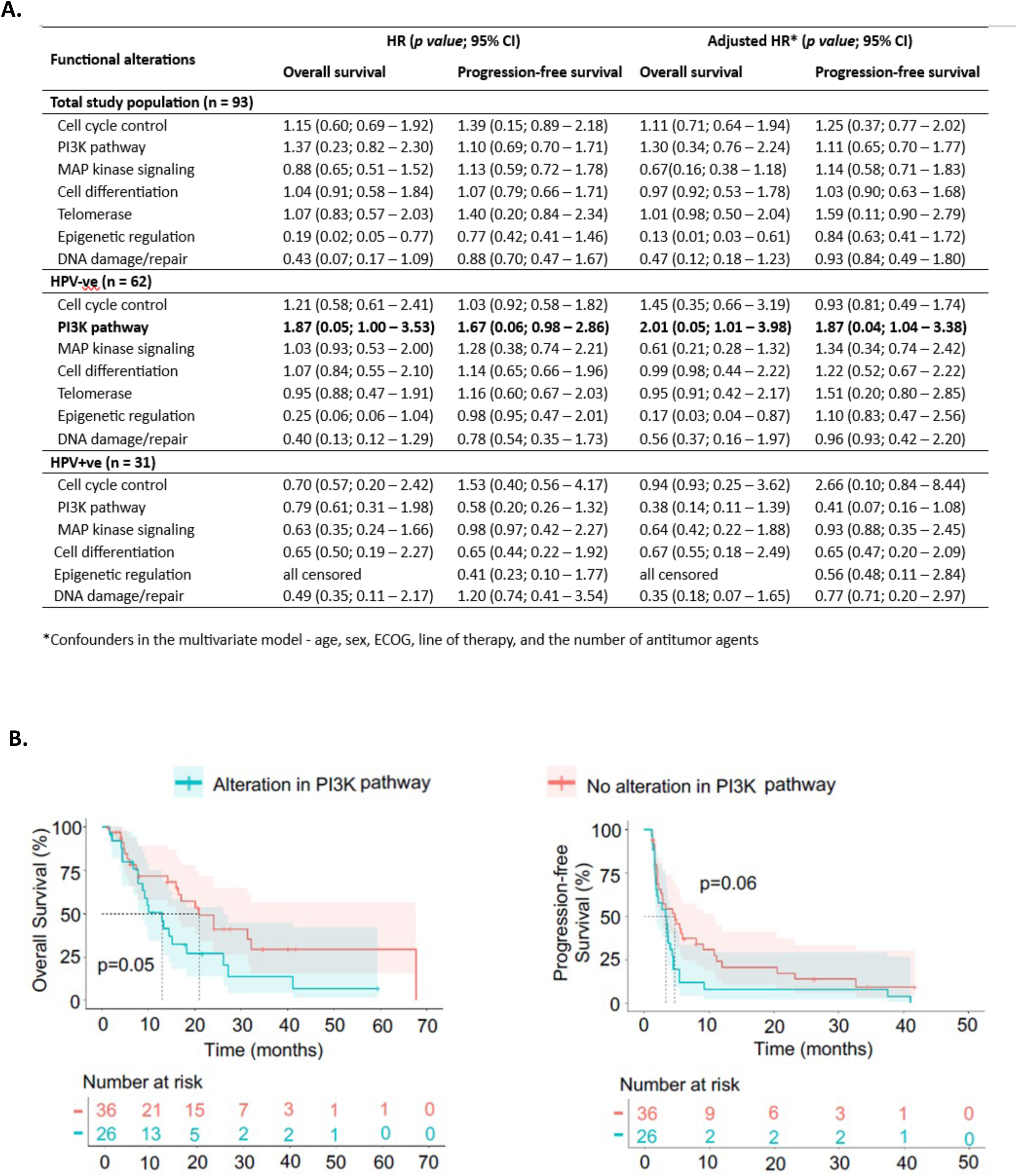

We examined the frequency and types of *PIK3CA* gene and its correlation with response to ICI in our cohort of 93 patients. We observed *PIK3CA* gene alterations, including mutations and amplifications, in 32.1% of non-responders compared to 27.2% of responders to immunotherapy (Figure 3A). In HPV-ve HNSCC patients, *PIK3CA* alterations were observed in 31.5% of non-responders and 9.1% of responders (Figure 3B). Detailed analysis of *PIK3CA* mutation patterns identified 13 different driving mutations, including E542K, E545K, and H1074Y (Figure 3C). Most of these mutations in *PIK3CA* are either known to be pathogenic or likely pathogenic in the helical and kinase domains based on the ClinVar analysis (Figure 3D) and these mutations were also reported in the TCGA study of HNSCC patients (Figure S2B). Additionally, *PIK3CA* amplification was observed in 7.1% of non-responders and 4.5% of responders (Figure 3E). Overall, we identified a higher frequency of mutations and amplifications in the *PIK3CA* gene among non-responders compared to responders to immunotherapy in HPV-ve HNSCC patients. While the role of *PIK3CA* activating mutations and amplification as drivers of oncogenic programs is well established, their influence on the tumor immune microenvironment and response to immunotherapy in patients, including patients with HNSCC has not been elucidated.

**Figure 3.**
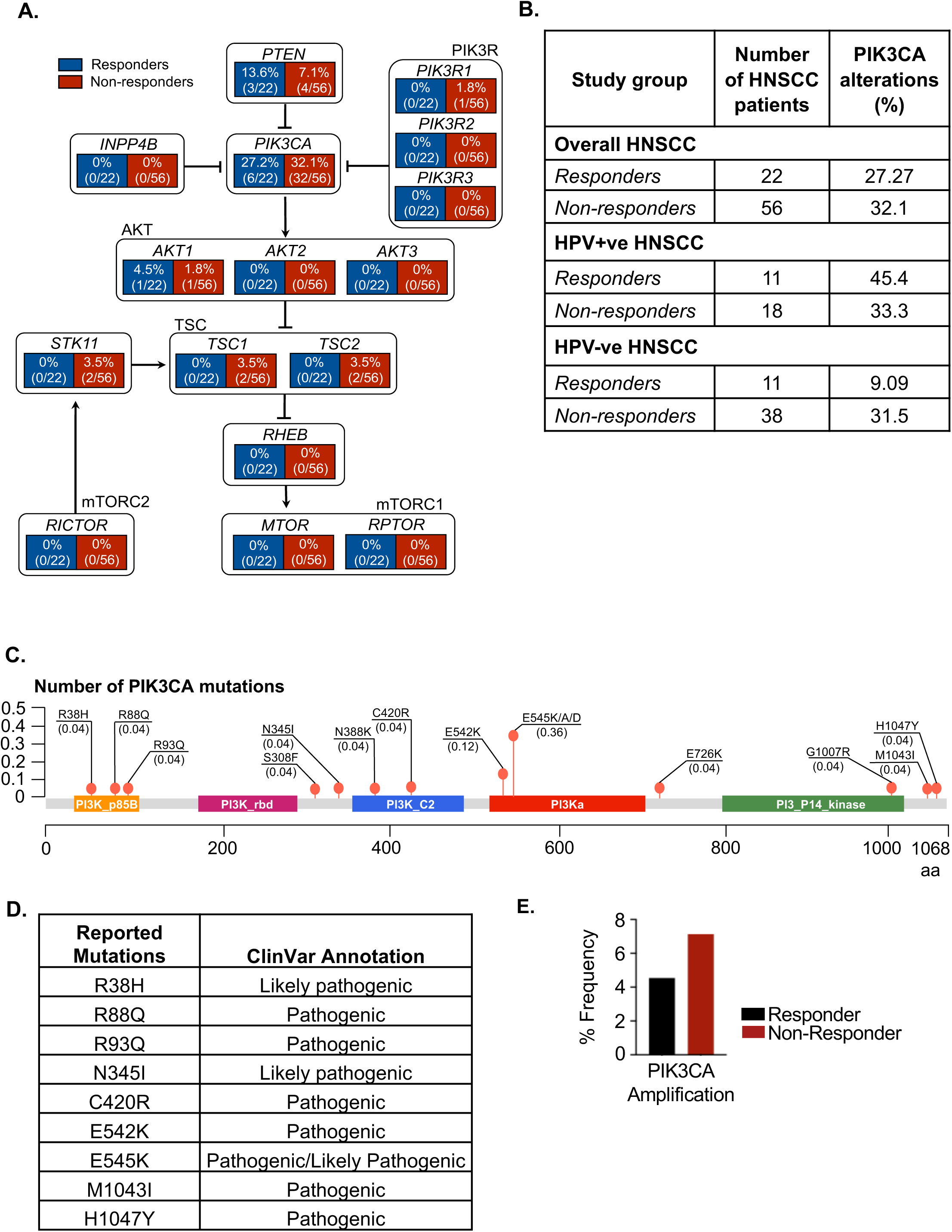

### Impact of *PIK3CA* on tumor immune microenvironment in HNSCC

Previous studies have shown that activated T-cell tumor infiltration is predictive of the response to ICI therapy^24–26^. To better understand the role of the PI3K pathway in regulating the immune response, we investigated if there was a correlation between *PIK3CA* gene expression and CD8+ T-cell tumor infiltration level utilizing the Tumor Immune Dysfunction and Inclusion (TIDE) approach. In our analysis of the Cancer Genome Atlas (TCGA) dataset utilizing the Tumor IMmune Estimation Resource (TIMER) algorithm, we identified an inverse correlation of the *PIK3CA* gene expression to CD8+ T-cell infiltration in all HNSCC and the HPV-ve HNSCC population but not in the HPV+ve group of patients (Figure S3A) (Figure 4A). Similarly, the gain of copy number and amplification of the *PIK3CA* gene were also associated with a significantly reduced number of tumor cytotoxic T-lymphocytes (CTL) in HNSCC patients overall (Figure S3B) and in HPV-ve HNSCC patients (Figure 4B).

**Figure 4.**
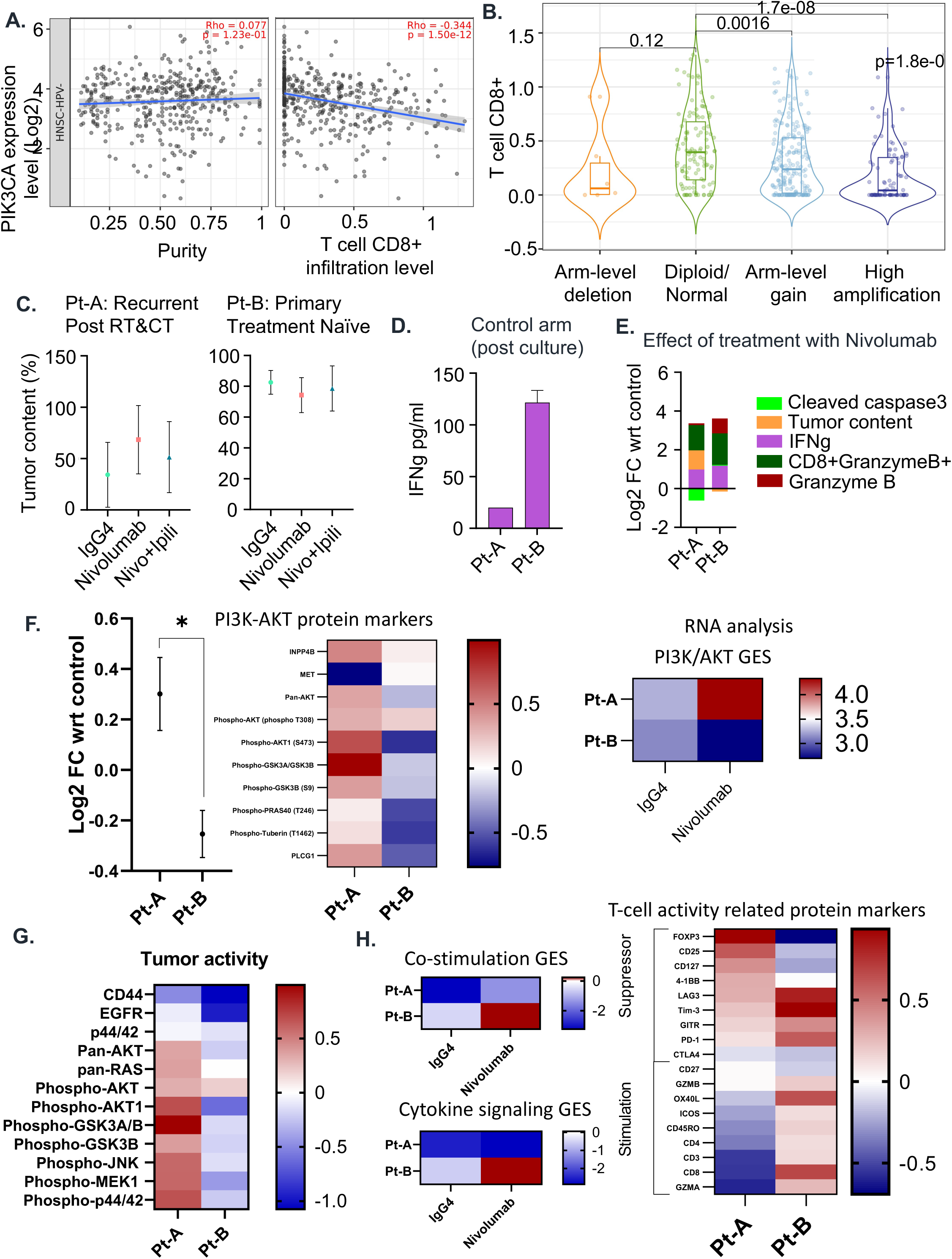
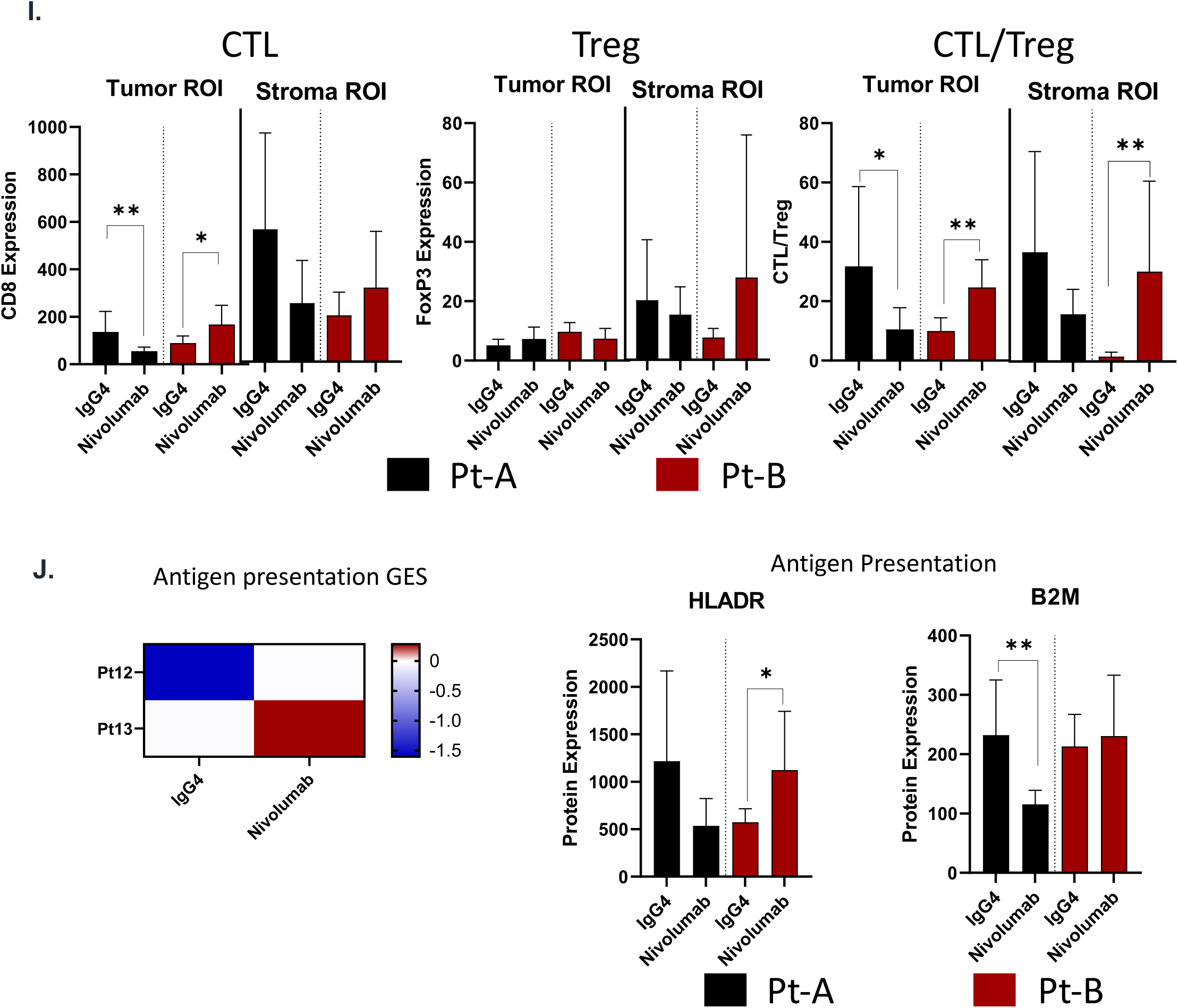

To evaluate the correlation between cell signaling pathways and responses to the ICI therapy, transcriptomic features were examined in two distinct HPV-ve HNSCC patient-derived tissues (Pt-A and Pt-B) using the Farcast^TM^ TruTumor platform. Pt-A had an aggressive locally advanced Stage IVA tumor which had recurred within the first year of definitive chemoradiation. Pt-B had a localized stage II primary tumor that had not undergone any treatment. Both tumors were treated either with nivolumab alone or in combination with ipilimumab (anti-cytotoxic T-lymphocyte-associated protein 4 (CTLA-4)). After nivolumab treatment with and without ipilimumab, Pt-A showed an increasing trend in tumor content with decreased caspase3 staining and morphologically preserved tumor cells compared to Pt-B (Figure 4C, E, Supp Figure S4B), indicating resistance phenotype in Pt-A.

Additionally, Pt-A demonstrated a lower level of interferon-gamma (IFNϒ) secretion over 72 hours in the absence of treatment compared to Pt-B, further suggesting a more immunosuppressive tumor phenotype of Pt-A compared to Pt-B (Figure 4D). While both patient samples showed a similar increase in the levels of IFNϒ secretion upon Nivolumab treatment, Pt-A exhibited a smaller increase in cytotoxic T-cell-mediated cytotoxicity (assessed by the prevalence of CD8+GranzymeB+ cells and granzyme B release) than Pt-B (Figure 4E).

Next, Pt-A and Pt-B’s gene and protein expression profiles were compared using the NanoString I/O 360 PanCancer panel and the 72-marker protein DSP GeoMx panel. While the mRNA expression data from the I/O 360 panel provided bulk transcriptomic information, DSP GeoMx assay provided spatial insights into the response phenotype. We observed the constitutive hyperactivation of PI3K pathway both at protein (in the tumor compartment, *p* = 0.04) and mRNA expression levels in Pt-A compared to Pt-B, upon nivolumab treatment (Figure 4F). Proteins associated with tumor progression were over-expressed in the tumor compartment (Figure 4G), with repression of proteins involved in T-cell activity (CD8, GZMA, GZMB, CD45RO, OX40L) and antigen presentation (HLA-DR) in Pt-A (Figure 4H). At the same time, immune suppressive markers such as FoxP3 within tumor increased in Pt-A compared to Pt-B post nivolumab treatment, suggesting that in Pt-A tumor there was increased immunosuppressive effect of regulatory T cells (Tregs) (Figure 4H). The expression pattern of these gene sets was the opposite in Pt-B with T-cell activity related genes over-expressed compared to those related to tumor progression (Figure 4F and G).

To correlate the observed phenotypes in the two samples with the spatial distribution of CD8+ CTL and Tregs, we quantified their spatial content in the tumor and stromal compartments, respectively. On treatment with nivolumab, recruitment of CTLs within the tumor compartment showed a significant decrease in Pt-A (*p* = 0.002) whereas a significant increase was observed in Pt-B (*p* = 0.02). The stromal compartment showed a similar but non-significant trend in both patients. On the other hand, FoxP3+ Tregs did not significantly change in tumor and stromal compartments of either samples, effectively leading to a significant increase CTL/Treg ratio in both tumor (*p* = 0.008) and stromal compartment (*p* = 0.001) for Pt-B. A significant decrease in CTL/Treg ratio was observed for Pt-A within the tumor compartment (*p* = 0.03) but remained unchanged in the stromal compartment (Figure 4I). To validate our findings, we analyzed if PIK3CA pathway contributes to predicting response to immunotherapy utilizing patient tumors that were cultured as histoculture. We found that patients with higher activity of PI3K pathway as assessed by the downstream target gene signature, exhibit a trend of poorer response to immunotherapy compared to patients with lower PI3K activity (Supp Figure S4C). On treatment with nivolumab, the expression of antigen presentation-related proteins, HLA-DR and B2M significantly decreased in Pt-A compared to control (*p* = 0.002). Nivolumab treatment induced a significant increase in HLA-DR (*p* = 0.02) but no change in B2M expression in the tumor compartment (Figure 4J).

### PIK3CA expression inversely correlates to Interferon-gamma responsive genes

Immune hot (or inflamed) tumor phenotype is characterized by the expression of immune response genes, including antigen processing machinery (APM), MHC class, interferon ϒ (IFNϒ) responsive genes, and proinflammatory chemokines that help in the recruitment of effector T-cells and predict a better response to immunotherapy^27,28^. Immune cold tumors are characterized by lower expression of immune response genes and impaired anti-tumor immune response. We further analyzed the TCGA data of 530 HNSCC patients to identify an association of *PIK3CA* gene expression with immune response genes. Gene expression analysis revealed a negative correlation between *PIK3CA* expression with multiple genes involved in APM, including *HLA-A, B, C, B2M, TAP1* and *TAP2,* and chemokine gene *CXCL10* (Figure 5A-D, S5B), consistent with the immune cold phenotype we observed with decreased infiltration of cytotoxic T cells. Furthermore, there was a trend of negative correlation between *PIK3CA* and the expression of STAT1, an essential IFNϒ responsive gene (Figure S5A). These results highlight that *PIK3CA* may negatively regulate the expression of pro-immune IFNϒ responsive and APM-associated genes, thereby impairing the activity and infiltrative capacity of CTLs in HNSCC patients.

**Figure 5.**
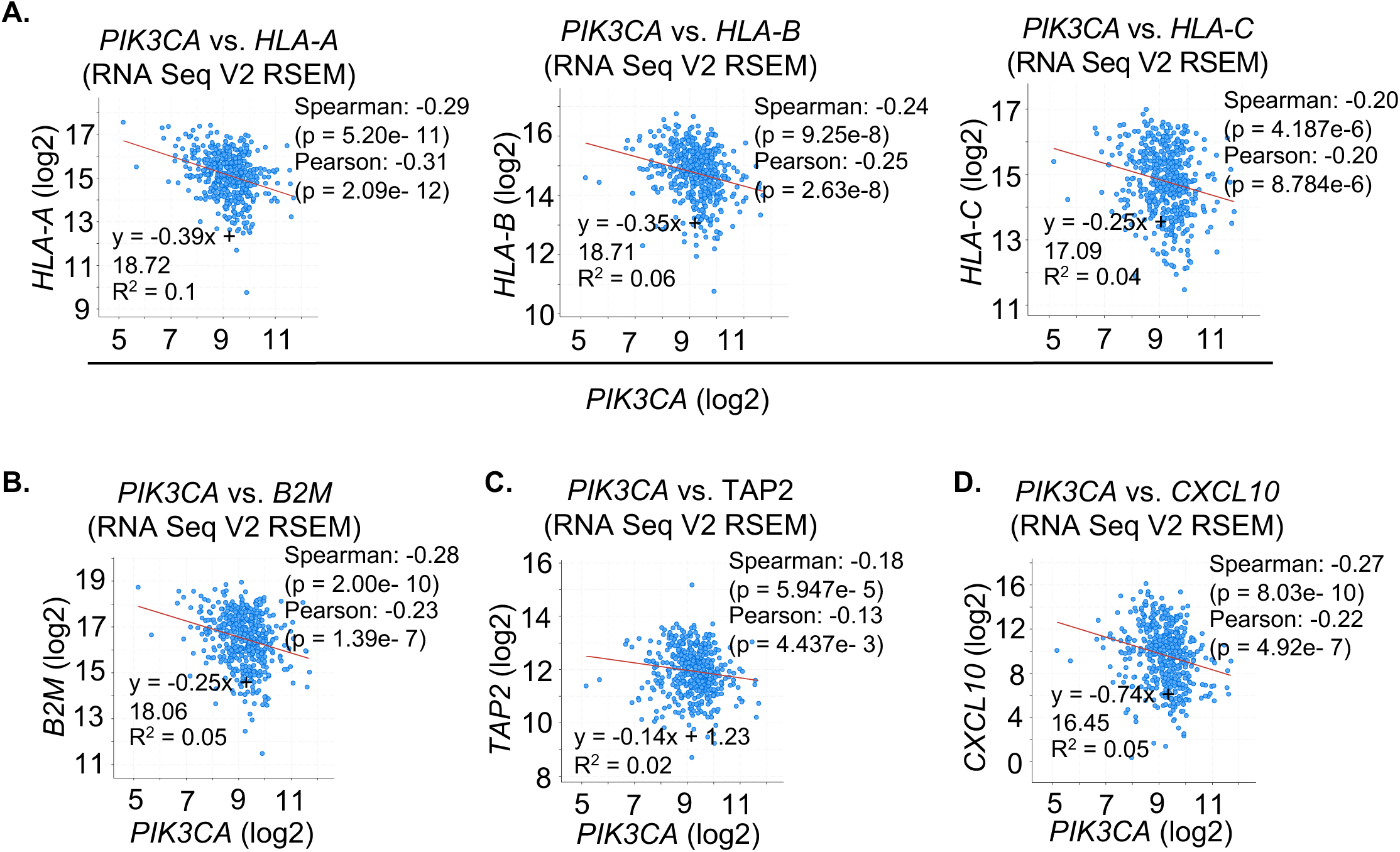

### *In vitro* inhibition of *PIK3CA* in HNSCC cell lines modifies immune gene expression

We tested the sensitivity of multiple HNSCC cell lines to BYL719 (alpelisib), a PI3K-alpha subunit specific inhibitor, and GDC-0941 (pictilisib), a pan-PI3K class I inhibitor utilizing the cell titer glo assay (Figure 6A). Among the cells we have utilized for studying the contribution of PIK3CA mutation to sensitivity to PI3K inhibitors, alpelisib and pictilisib, HSC4, CAL33, DETROIT562, and HSC2 cells are *PIK3CA-*mutant and BICR22, BHY, CAL27, and FaDu cells are *PIK3CA w*ild-type HNSCC cells (Figure 6B). As expected, *PIK3CA-*mutant cell lines demonstrated higher sensitivity to both alpelisib and pictilisib with a lower IC50 value compared to the wild-type cell lines suggesting the dependency of the cells on PI3K pathway (Figures 6C and 6D). Furthermore, the impact of PI3K inhibitors on immune gene expression was examined in *PIK3CA*-mutant CAL33 (H1047R) and HSC4 (E545K) cell lines, as well as the PIK3CA wild-type FaDu cell line. Treatment with alpelisib induced a significant increase in crucial immune response gene expressions such as HLA, APM, IFNϒ and chemokine genes in the CAL33 and HSC4 cells (Figure 7A), but not in FaDu cells (Figure 7A). Similarly, pictilisib treatment induced a significant increase of immune response genes in CAL33 and HSC4 cells, but not in FaDu cells (Figure 7A).

**Figure 6.**
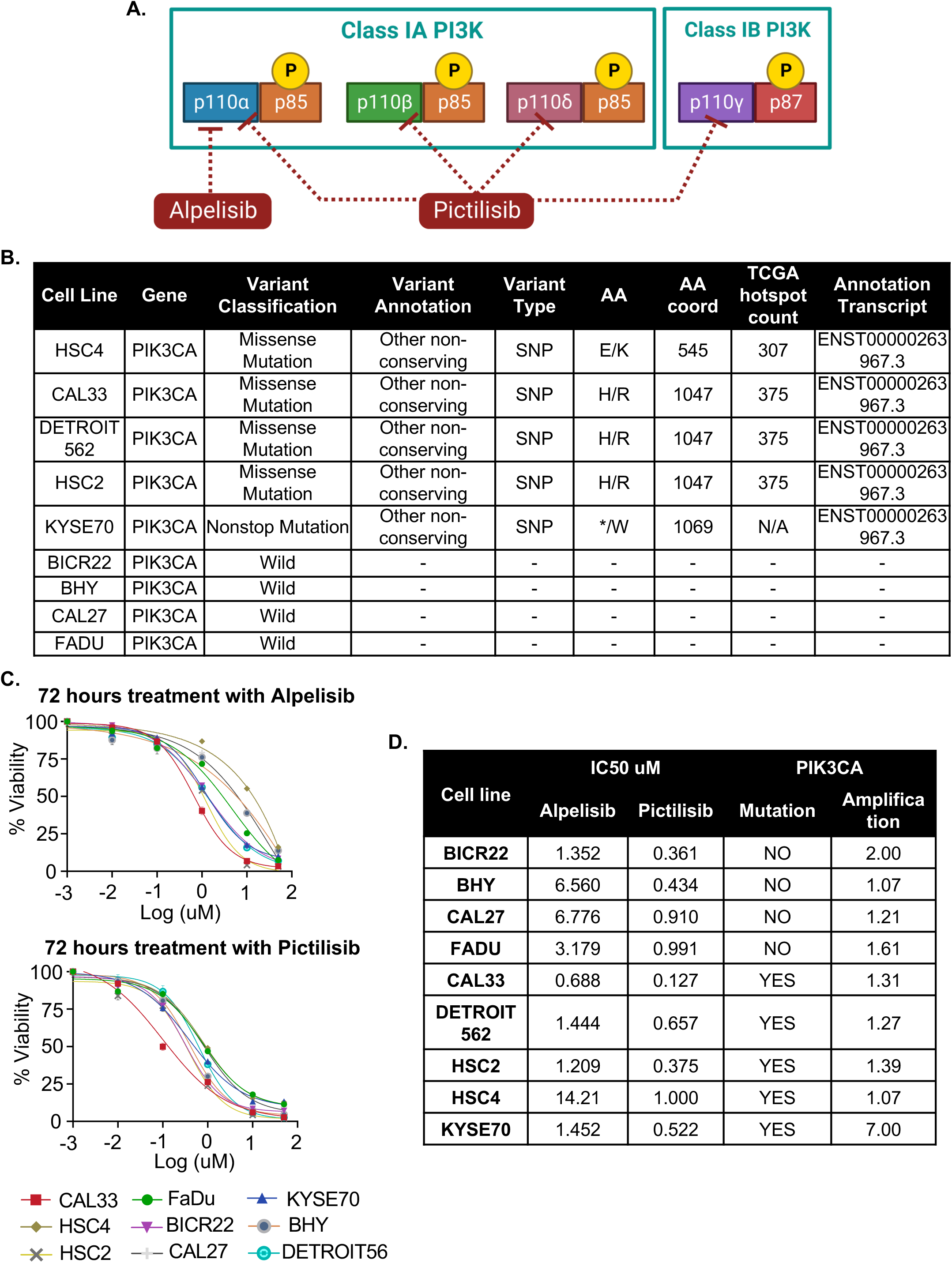

**Figure 7.**
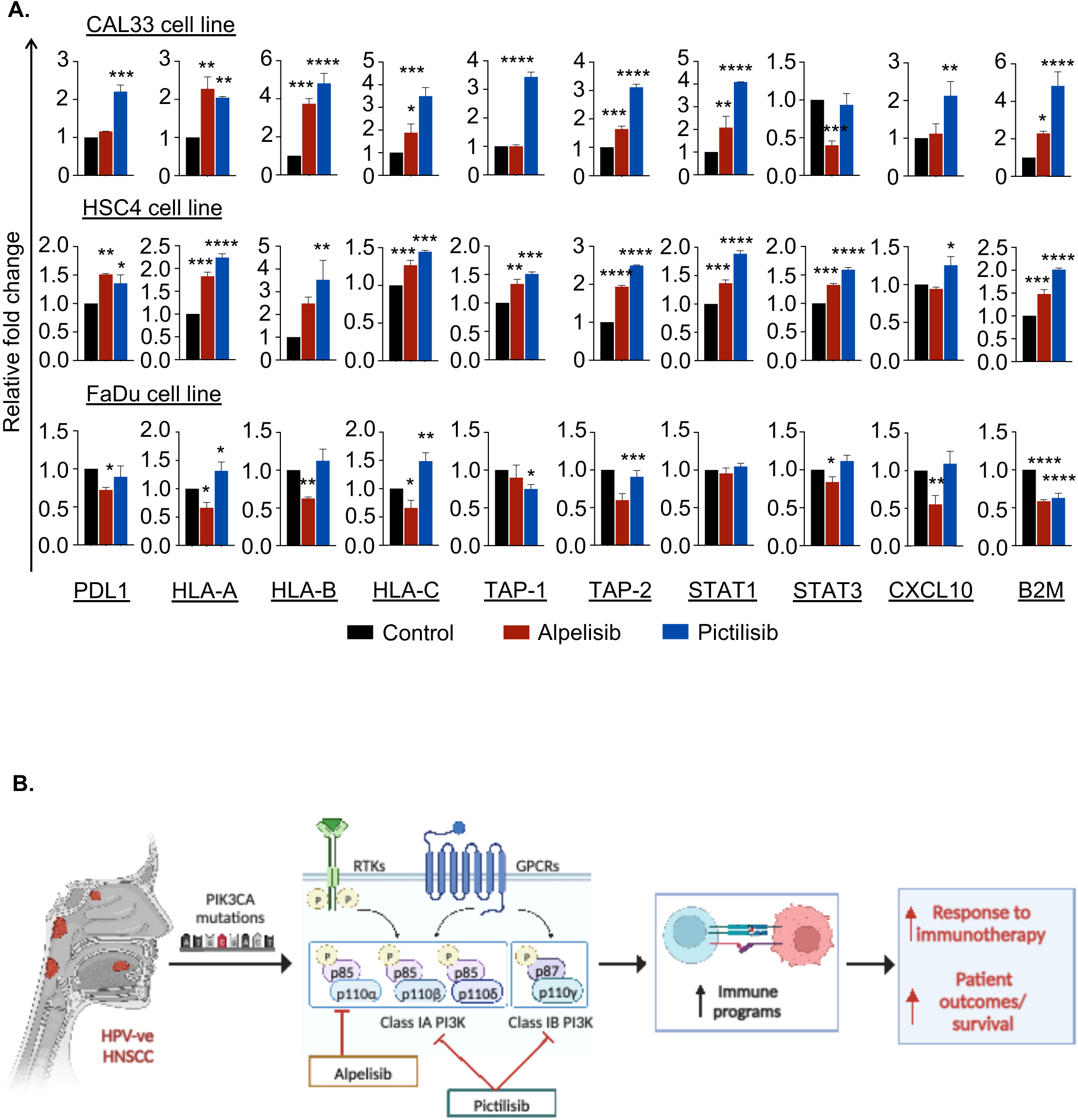

## DISCUSSION

Our retrospective analysis demonstrated that the patients with R/M HNSCC whose tumor harbored PI3K signaling pathway alterations had inferior survival and response to immunotherapy. This association seems to be limited to HPV-ve population and suggests that the PI3K signaling pathway may contribute to immune evasion in this population. The prognostic implication of *PIK3CA* mutation in HNSCC has not been clearly established, and published data provides conflicting conclusions^31,32^. Cochicho et al reported a case series demonstrating a superior survival in patients with *PIK3CA-*mutant HNSCC compared to the wild-type in HPV+ve patients, but numerically inferior survival in HPV-ve population^32^. However, the post hoc analysis of de-intensified chemoradiation trials (NCT02281955, NCT03077243) suggested a worse disease-free survival in patients with *PIK3CA-*mutant HPV+ve locally advanced oropharyngeal cancer^31^. The data of prognostic value of PI3K pathway alteration in HPV-ve population are very limited. Our single-institution retrospective patient outcome data is a retrospective analysis and the small sample size with uncontrolled diverse ICI regimens received among different cohorts. The radiologic response assessment was based on the treating clinicians’ assessments and radiology reports and not based on the Response Evaluation Criteria in Solid Tumors (RECIST) criteria. Thus, our data reports the first observation of alterations of PIK3CA as a key prognostic factor in HPV-ve HNSCC patients who received ICI.

PI3K pathway regulates multiple biological processes including DNA repair, glucose homeostasis, cell proliferation and survival, and protein synthesis^33–36^. PI3K comprises of two subunits, regulatory (p85) and catalytic (p110) subunits. In quiescent cells, the regulatory unit inhibits the catalytic subunit activity^37^. *PIK3CA* gene encodes for the class IA p110α subunit. A wide array of *PIK3CA* mutations have been identified in various solid tumors with 3 most frequent hotspots of which E542 and E545 are in the helical domain, and H1047 is in the kinase domain^13,38,39^. The high frequency of *PIK3CA* mutations in HNSCC has been highlighted in various studies, and shown to be enriched in HPV+ve HNSCC patients^40–45^. The TCGA reported *PIK3CA* gene mutation or amplification in 56% of patients with HPV+ve HNSCC, and 34% in HPV-ve HNSCC patients^43^. The E545 is most commonly mutated followed by the E542 region, and mutations in the H1047 hotspot is rare in HNSCC^32,46,47^. In our patient cohort who received ICI, mutations or amplification of PIK3CA were observed in 27% of patients, 36% which were observed in HPV+ve and 23% in HPV-ve, respectively, with the E545 alteration being the most common mutation reported in TCGA for HNSCC patients. *PIK3CA* alteration was mutually exclusive with other PI3K pathway gene alterations such as *PTEN, AKT1,* and *PIK3R*.

Several PI3K inhibitors have demonstrated antitumor efficacy and are approved for treatment for *PIK3CA-*mutant advanced breast cancer. Alpelisib is an alpha subunit-selective PI3K inhibitor that was approved by the U.S. Food and Drug Administration (FDA) in combination with Fulvestrant for the treatment of advanced *PIK3CA*-mutant breast cancer^48^. Alpelisib has previously demonstrated pre-clinical anti-tumor activity in *PIK3CA-*mutant HNSCC models but has limited efficacy as monotherapy in early phase clinical trials^29,30^. Pictilisib is a pan Class 1 PI3K inhibitor with activity against all 4 isoforms (alpha, beta, gamma, and delta). Pictilisib in combination with paclitaxel was shown to have antitumor activity in breast cancer patients^50^. While single agent activity was limited, chemotherapy and cetuximab combination data were encouraging^51–54^. But the toxicity rates were high with combinatory approaches^53,54^. We observed selective sensitivity to alpelisib and pictilisib in *PIK3CA-*mutant versus *PIK3CA* wild-type HPV-ve HNSCC cells. Treatment with both alpelisib and pictilisib increased expression of immune response, antigen presentation, and chemokine genes in *PIK3CA*-mutant HNSCC cell lines but not in *PIK3CA* wild-type cell lines. Based on these findings, we propose a model indicating utilization of inhibitors against PI3K to reactivate the immune programs. (Figure 7B).

The potential impact of PI3K pathway on tumor immune microenvironment and immune phenotypic changes have been reported in solid tumors^55–59^. Interestingly, the impact of PI3K pathway alteration on the tumor immune microenvironment appears conflicting in different tumor types, suggesting the impact may be tumor-type dependent. Early studies reported by Chandrasekaran et at utilizing PI3K inhibition showed enhanced IFNϒ-mediated induction of MHC molecules and STAT1^60^ in a panel of cell lines. PI3K-mediated alterations in MHC class I molecules have been shown to impact T-cell infiltration into tumors, including pancreatic ductal adenocarcinoma mouse model and few cell line models including HPV+ve HNSCC cell line and a melanoma cell line^56,61^. High mTORC1 activity has been associated with increased CD276 expression and immunosuppressive phenotypes in the TCGA cohort^62^. Targeting CD276 in mTORC1 high cells was shown to increase MHC class II expression leading to anti-tumor immunity. Wang et al showed that PI3K pathway is activated in *PIK3CA* wild-type HNSCC through the HER3 receptor tyrosine kinase, and downstream activation of this pathway induced immune suppressive cytokines and recruitment of immune suppressive cells in the tumor microenvironment^63^. Blocking HER3 with a monoclonal antibody, CDX3379, reversed such immune suppressive phenotypic changes and synergized with anti-PD1 anti-tumor activity. However, the data on the role of *PIK3CA* alteration and PI3K pathway dysregulation in the context of immunotherapy response in HNSCC patients has not been reported.

In summary, we identified that alterations in PIK3CA gene and other components of the PI3K pathway correlates to poorer response to ICI therapy in HPV-ve HNSCC patients. Alterations in PI3K pathway mediate resistance to immunotherapy agents in patients through remodeling of immune gene expression, thereby promoting immune evasion. We show that pharmacologic PI3K inhibition reactivates APM, MHC, and IFNϒ responsive genes thus priming the tumor cells to immunotherapy in the context of HPV-ve HNSCC. Our findings support a potential strategy of combining PI3K inhibitors with ICIs to improve outcomes in HPV-ve HNSCC patients and paves the way for establishing clinical trials following identification of mutations in PIK3CA gene supporting the idea of personalized medicine (Figure 7B). This study paves way for a larger group and multi-institutional study to conduct a clinical trial targeting the PI3K pathway components to improve the response and/or overcome the resistance to ICI.

## Materials and Methods

### Patient cohort

We performed a retrospective search of the electronic medical record at Massachusetts General Cancer Center to identify patients with R/M HNSCC who underwent systemic therapy with ICI from 2016 to 2022 and met the following criteria: age ≥18; histologically confirmed squamous cell carcinoma of the head and neck; received palliative ICI either monotherapy or in combination with other immune-based agents excluding chemotherapy combination; available next-generation sequencing-based molecular test results. Clinicopathologic features, treatment history, and outcomes were captured. The primary outcome measure of interest was OS. Additionally, PFS and radiographic response were analyzed. Of note, we define the radiographic response as unequivocal decrease of overall tumor size(s) regardless of the magnitude. The Response Evaluation Criteria in Solid Tumors (RECIST) was not used for this retrospective analysis as the staging scans in analyzed population were not universally evaluated using the strict RECIST criteria and implementation of independent radiology review was not feasible. This study was conducted under the IRB Protocol no. 2018P001456.

### Statistical analysis

We applied Cox proportional hazard model to elucidate whether functional alterations affect overall survival and progression-free survival in HPV-ve and HPV+ve patients. Univariate and multivariate analyses were used to calculate hazard ratios and 95% confidence intervals. Confounders in the multivariate model - age, sex, ECOG, line of therapy, and the number of antitumor agents - were selected based on the previous studies and clinical significance. The proportional hazard assumption for the main results was checked using Schoenfeld residuals to ensure the validity of the models. Kaplan-Meier curves stratified by the alteration status of PI3K pathway in HPV-ve patients were also demonstrated, with *p-value* derived by log rank test. The data were analyzed using the statistical software R, version of 4.3.0.

### Patient tissue samples and human tumor histoculture and multimodal assay workflow

Fresh surgically resected HPV-ve HNSCC tissue samples along with matched blood were collected from consented patients using a protocol (Protocol no. FCB-Protocol-01_v1_20Feb2021) approved by institutional ethics committee (approval no. VIEC/2021/APP/021). The tumor samples were processed to generate thin explants without enzymatic digestion retaining the tumor microenvironment. Blood was processed to separate plasma. The tumor explants were cultured with media and autologous plasma using the Farcast^TM^ TruTumor human tumor histoculture platform. The explants were treated either with 132 µg/ml Nivolumab (anti-PD1) or a combination of 132 µg/ml Nivolumab and 90.8 µg/ml Ipilimumab (anti-CTLA4) for 72 hrs. Fresh media containing drug was replaced every 24 hrs and the supernatant was collected and assayed for cytokine release. The tissue culture was terminated at 72 hrs and either fixed in Neutral Buffer Formalin (NBF) and converted into FFPE blocks or used to generate single cells for flowcytometry analysis.

### Histopathology and Immunohistochemistry

Four-micron sections containing slides from FFPE blocks were stained with Hematoxyllin and Eosin and tumor content was evaluated by certified pathologist. Cleaved caspase3 (PP 229 AA; RTU; Biocare) IHC was performed and scored by pathologist to determine tumor apoptosis.

### Cytokine release assay and Flow cytometry

We evaluated the IFNϒ and GranzymeB cytokine secretion in the collected culture supernatant using Luminex based cytokine analysis (Procarta Plex, Thermo Fisher Scientific). CD8 activity was determined by flowcytometry by evaluating CD8+GranzymeB+ cells.

### NanoString GeoMx DSP

Slides containing four-micron sections were used for NanoString DSP GeoMx. A 72-protein marker assay panel using panCK and DNA as morphological markers was performed. The proteins encompassed various pathways related to tumor and immune function (Details included in additional data section). panCK was used to segment tumor and stromal compartments. The data generated was analyzed by GeoMx DSP software. Three housekeeping proteins (GAPDH, Histone H3 & Ribosomal protein S6) were used to normalize the data. The normalized data for the selected Region of Interest (ROI) was used for marker expression analysis. We performed differential expression analysis for all the protein markers. Spatial analysis for CD8 and FoxP3 protein markers representing CTL and FoxP3 in tumor and stromal segments was performed.

### NanoString Gene expression Analysis

The RNA extracted arm-wise (control and nivolumab) from 8-10 four-micron sections Tissue Micro Array (TMA) FFPE block and tested using the NanoString PanCancer I/O 360 panel. RNA was quantified using Tape Station. 30-50ng of RNA based on DV200 concentration was used for running on the NanoString IO 360 panel. Data was normalized and analyzed using nSolver™ Data Analysis software.

### Mutation analysis

The patients in the study cohort were categorized into responders and non-responders based on their response to immunotherapy. The NGS was performed to analyze the mutation profile of the patients. The snapshot (oncoplot) of the mutations observed in individuals was prepared. The top 5 most altered genes namely, *TP53, PIK3CA, CDKN2A, TERT,* and *NOTCH* were selected for the comparative and functional analysis. The comparison of mutation frequency was performed between responders and non-responders and presented in the form of lollipops on the protein/gene graphs based on their amino acid or nucleotide position where the height of the lollipop represents the frequency of mutation in responder or non-responder.

The details of the mutations, namely, location, nucleotide change, change in amino acid, clinical significance, and functional consequence, were extracted from Ensembl database^64^ and presented in a descriptive table. For the functional analysis of genetic variations, we utilized 3 different algorithms/tools namely, SIFT^65^, POLYPHEN^66^, and Mutation Assessor (http://mutationassesor.org/r2/; assessed as on 25 Jan 2023). SIFT score is used to assess the tolerance of the protein towards the change in amino acid sequence due to the mutation present in the gene^65^. POLYPHEN score suggests the tolerance of the change in amino acid sequence across various polymorphic phenotypes based on an evolutionary comparison^66^. The mutation assessor score predicts the functional impact of the amino acid change by comparing different protein homologs (http://mutationassesor.org/r2/; assessed as on 25 Jan 2023). The utilization of 3 different algorithms namely, SIFT, POLYPHEN, and Mutation Assessor helped in the selection of the mutations that had the most lethal or deleterious impact on the protein function. A concordance analysis was performed to select the most lethal mutations unanimously predicted as deleterious by all 3 algorithms.

### Cell viability assay

The cell viability assay was performed to determine the IC50 for drug treatment as described earlier^67^. Briefly, HNSCC cells were seeded in 96 well plates and treated with different concentrations of drugs. The viability of cells was measured using CellTiter-Glo” Luminescent assay (Cat #: G7570, Promega) after 72 hours of drug treatment. A microplate reader (SpectraMax M5, Molecular Devices, Sunnyvale, CA, USA) was used for measuring the luminescence. The dose-response curve and estimation of IC50 were performed using GraphPad Prism 7.0.

### RNA extraction and qPCR

Total RNA was extracted from cells treated with Alpelisib and Pictilisib drugs using the Trizol method as described earlier^67^. The RNA quality and quantity were assessed using Nanodrop (Thermo Fisher Scientific) and 1μg of total RNA was used for cDNA synthesis with Applied Biosystems^TM^ High-Capacity cDNA Reverse Transcription Kit with RNase Inhibitor (Cat #: 4374967, Thermo Fisher Scientific). The gene expression analysis was performed using Power SYBR^TM^ Green PCR Master Mix (Thermo Fisher Scientific, Cat #: 4368702) according to the manufacturer’s instructions. The expression of genes was normalized to 18S house-keeping gene expression in each sample. The data were represented as a fold change of gene expression in drug-treated cells in comparison to control cells. The primer sequences used for analyzing the gene expression are presented in the supplementary table (table S3).

## Acknowledgements and Declaration of Interests

LW receives honoraria for advisory roles from Bayer HealthCare Pharmaceuticals, Blueprint Pharmaceuticals, Coherus, Curie Therapeutics, Eli Lilly, Eisai, EMD Serono, Exelixis, Genentech USA, Merck, Morphic Therapeutics, Tome Biosciences and data safety monitoring board for PDS Biotechnology Corporation. JCP serves on the Scientific Advisory Boards, Consultant, or Expert Witness for GC Cell, Selecxine, Hanmi Pharmaceutical, and on the clinical Research Support/Data Safety Monitoring Board for MitoImmune. We thank all the members of Saladi lab for helpful comments and scientific discussions. NPB and SS are employees of Farcast Biosciences. This work is funded by Mike Toth Head and Neck cancer center funds to SVS. SVS is cofounder of a stealth mode startup not related to the work in this manuscript.

