## Supplemental data and figures for "Impact of PI3K pathway alterations on response to immune checkpoint inhibitors in HPV-negative head and neck squamous cell carcinoma"

**Figure S1.**

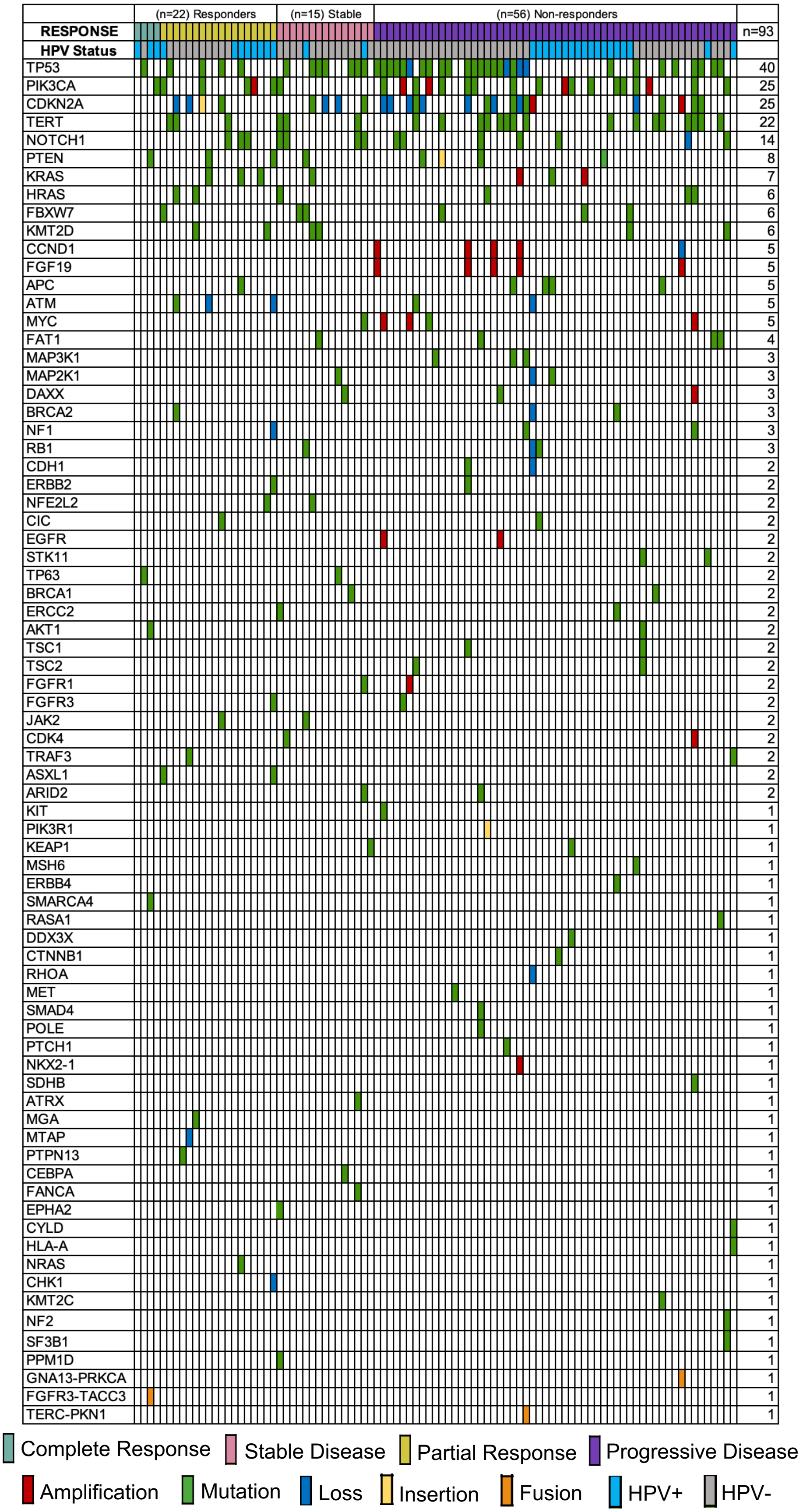


**Figure S1. Overview of HPV and immunotherapy response status and mutation signature analysis in HSNCC patients.** 93 samples of HNSCC patients with metastatic/recurrent disease undergoing molecular pathologic diagnostic before starting immunotherapy treatment. All the genetic alterations including mutations and amplifications are plotted. Out of 93 patients, 22 patients responded to immune checkpoint inhibitors (including 4 patients with complete response and 19 patients with partial response), 15 patients had stable disease and 56 progressed in the disease course despite treatment.

**Figure S2.**

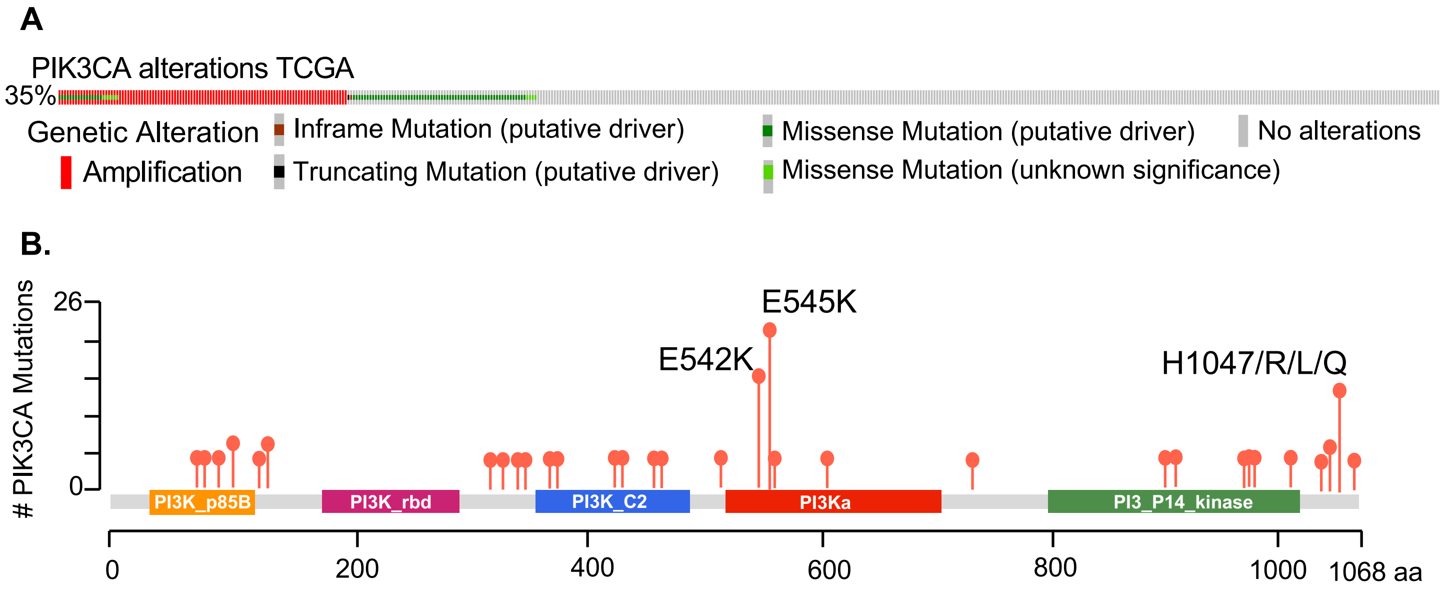
A
PIK3CA alterations TCGA
35%
Genetic Alteration
Amplification
Inframe Mutation (putative driver)
Truncating Mutation (putative driver)
Missense Mutation (putative driver)
Missense Mutation (unknown significance)
No alterations
B.
### PIK3CA Mutations
26
0
E545K
E542K
H1047/R/L/Q
PI3K_p85B
PI3K_rbd
PI3K_C2
PI3Ka
PI3_PI4_kinase
0
200
400
600
800
1000
1068 aa

**Figure S3.**

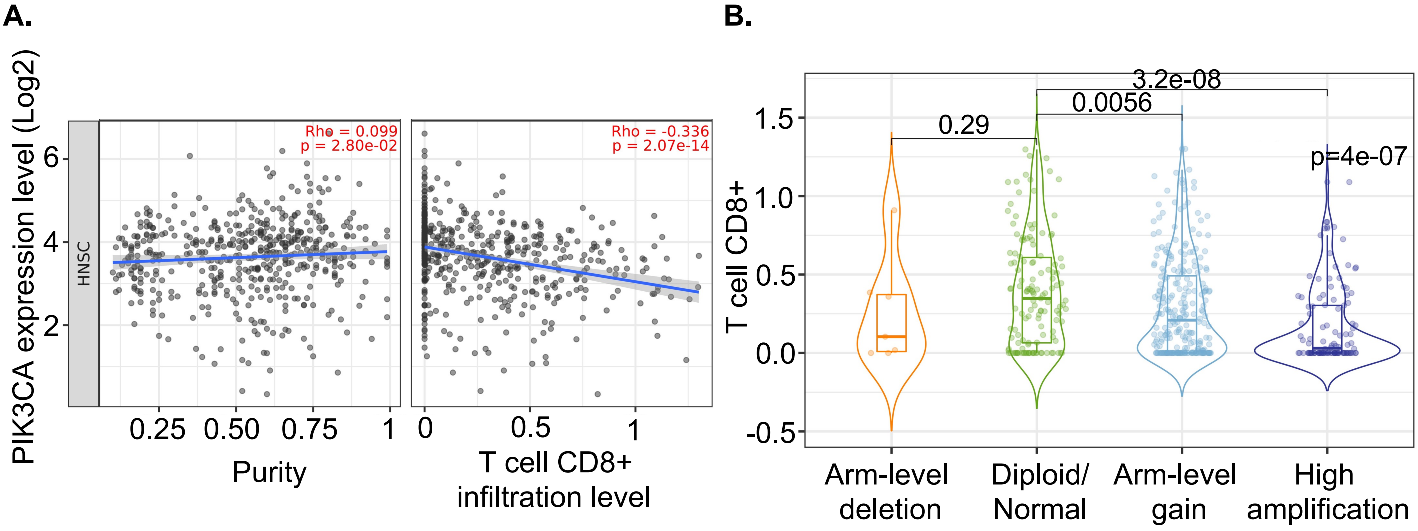


**Figure S3. PIK3CA gene expression and copy number correlates with CD8+ T lymphocyte in HNSCC patients from TCGA dataset. A.** The correlation between PIK3CA expression and T cell CD8+ infiltration level. **B.** Differential T cell CD8+ infiltration level in HNSCC HPV-negative patients. P value < 0.05 was considered statistically significant.

**Figure S4.**

**A.**

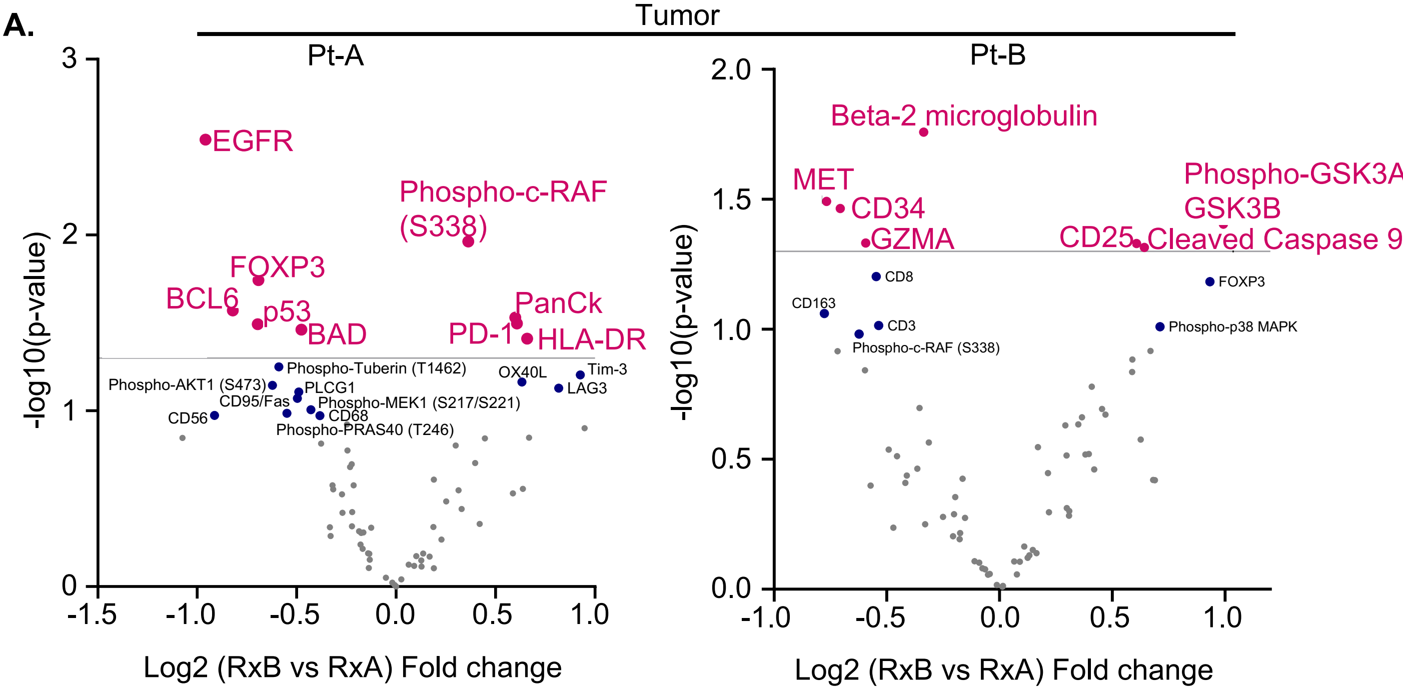


**B.**

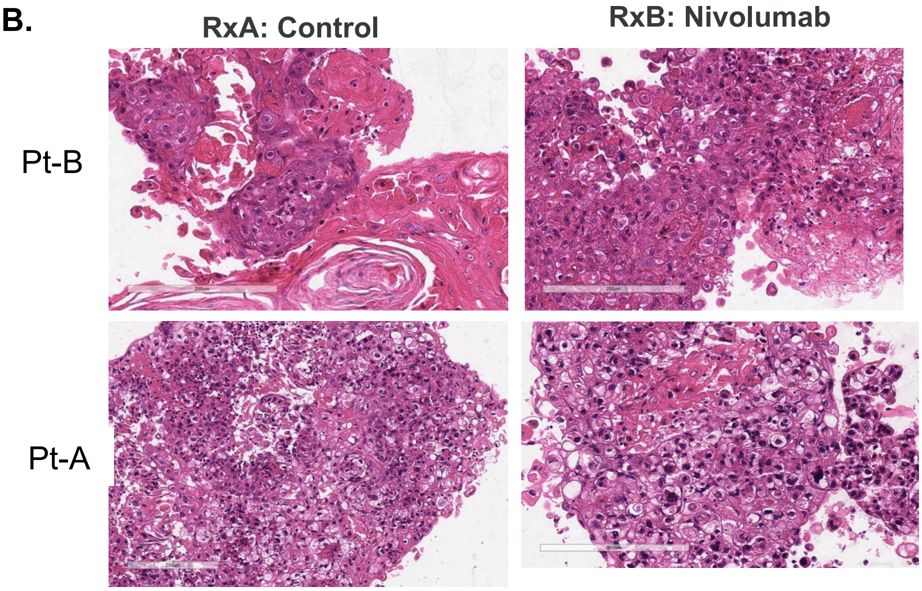


**C.**

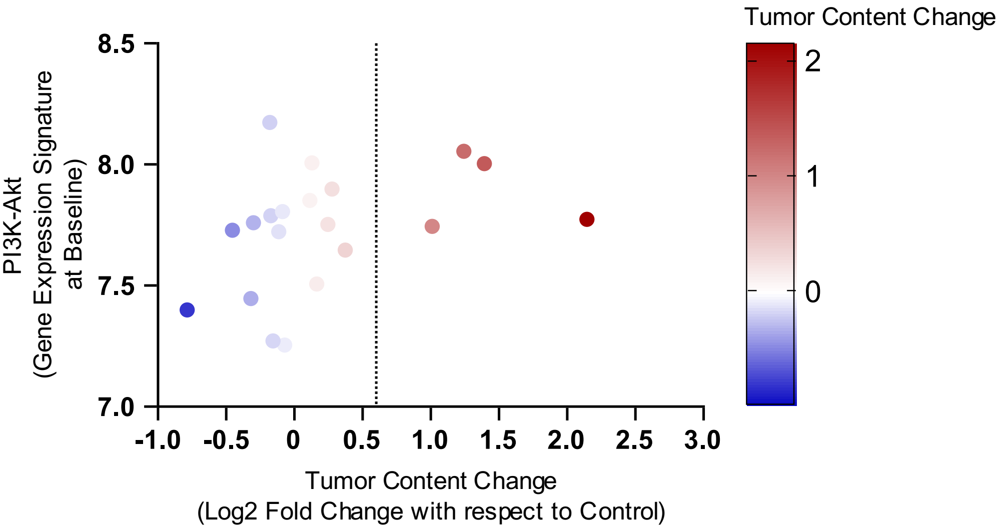

**Figure S5. Correlation of PIK3CA to immune genes.** PIK3CA expression shows inverse correlation to the immune genes **A.** STAT1. **B.** TAP1.

**A.**

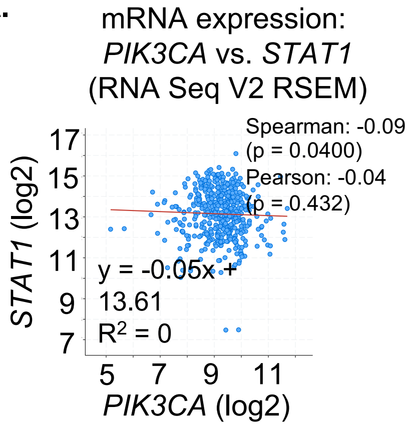


**B.**

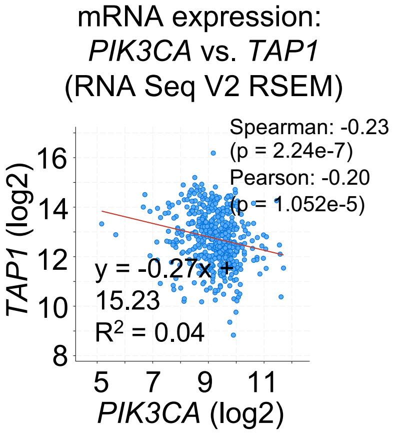

**Supplemental Table 1.** Baseline characteristics of the study population.

| Characteristics | Total (N = 93) |
|---|---|
| **Age, Median (Range)** | 65 (29 – 90) |
| **Sex, N (%)** | |
| Male | 72 (77.4) |
| Female | 21 (22.6) |
| **Race, N (%)** | |
| White | 80 (86.0) |
| Asian | 12 (12.9) |
| Black | 1 (1.1) |
| **The anatomical site, N (%)** | |
| Oropharynx | 33 (35.5) |
| Oral cavity | 26 (28.0) |
| Nasopharynx | 12 (12.9) |
| Larynx/hypopharynx | 13 (14.0) |
| Sinus/nasal cavity | 6 (6.5) |
| Unknown | 3 (3.2) |
| **ECOG status, N (%)** | |
| Good (≤ 1) | 78 (83.9) |
| Poor (≥ 2) | 15 (16.1) |
| **Smoking, N (%)** | |
| Yes* | 42 (45.2) |
| No | 51 (54.8) |
| **HPV status** | |
| Positive | 31 (33.3) |
| Negative | 62 (66.7) |
| **Immunotherapy agent†** | |
| Anti-PD-1 monotherapy | 74 (79.6) |
| Anti-PD-1 in combination | 19 (20.4) |
| **Line of immunotherapy** | |
| 1st line | 55 (59.1) |
| ≥ 2nd line | 38 (40.9) |
| **Follow-up time (months), Median (Range)** | 14.2 (1.4 – 67.6) |
| **CPS‡ (%), Median (Range)** | 10 (0 – 100) |

Abbreviation: N, number of people; ECOG, Eastern Cooperative Oncology Group; HPV, human papillomavirus; PD-1, program death-1; CPS, combined positive score

* Equal or greater than 10 pack years was defined as 'Yes' for the smoking variable.

† Subjects who were treated with a chemotherapy combination were excluded from the study population.

‡ CPS was available in 55.9% (N = 52) of total patients.

**Supplemental Table 2.** Prevalent (N > 10%) single genetic alterations in the study population.

| Genetic alteration | N (%) |
|---|---|
| **Total study population (N = 93)** | |
| TP53 | 40 (43.0) |
| CDKN2A | 25 (26.9) |
| PIK3CA | 25 (26.9) |
| TERT promoter | 23 (24.7) |
| NOTCH1 | 14 (15.1) |
| H/K/NRAS | 13 (14.0) |
| **HPV- (N = 62)** | |
| TP53 | 38 (61.3) |
| CDKN2A | 24 (38.7) |
| TERT promoter | 22 (35.5) |
| PIK3CA | 14 (22.6) |
| NOTCH1 | 11 (17.7) |
| H/K/NRAS | 8 (12.9) |
| **HPV+ (N = 31)** | |
| PIK3CA | 11 (35.5) |
| H/K/NRAS | 5 (16.1) |
| PTEN | 5 (16.1) |
| FBXW7 | 4 (12.9) |

Abbreviation: N, number of people

**Supplement Table 3.** Altered genetic function in the study population.

| Functional alteration | N (%) | | |
|---|---|---|---|
| | Total | HPV- | HPV+ |
| **The number of patients** | **93** | **62** | **31** |
| Cell cycle control | 49 (52.7) | 43 (69.4) | 6 (19.4) |
| PI3K/AKT/mTOR pathway | 41 (44.1) | 26 (41.9) | 15 (48.4) |
| MAP kinase signaling | 35 (37.6) | 25 (40.3) | 10 (32.3) |
| Cell differentiation | 30 (32.3) | 23 (37.1) | 7 (22.6) |
| Telomerase | 23 (24.7) | 22 (35.5) | 1 (3.2) |
| Chromatin remodeling/DNA methylation | 15 (16.1) | 10 (16.1) | 5 (16.1) |
| DNA damage/repair | 12 (12.9) | 8 (12.9) | 4 (12.9) |
| Wnt/β-catenin pathway | 6 (6.5) | 2 (3.2) | 4 (12.9) |
| Oxidative stress | 4 (4.3) | 3 (4.8) | 1 (3.2) |
| JAK/STAT pathway | 2 (2.2) | 1 (1.6) | 1 (3.2) |
| Others | 11 (11.8) | 9 (14.5) | 2 (6.5) |

Abbreviation: N, number of people

**Supplemental Table 4.** Primer sequences used for gene expression analysis.

| qPCR Oligos | Sequence |
|---|---|
| PDL1 F | ATGGTGGTGCCGACTACAA |
| PDL1 R | TCCAGATGACTTCGGCCTT |
| HLA-A F | GGCCCTGACCCAGACCTG |
| HLA-A R | GCACGAACTGCGTGTCGTC |
| HLA-B F | CATCGTGGGCATTGTTGCTG |
| HLA-B R | ACGCAGCCTGAGAGTAGC |
| HLA-C F | CTGGCCCTGACCGAGACCTG |
| HLA-C R | CGCTTGTACTTCTGTGTCTCC |
| TAP1 F | CCTGTGGCACAAACTCGGG |
| TAP1 R | ATCTCCCCAAGAGAGGAGAGGA |
| TAP2 F | TCGACTCACCCTCCTTTCTC |
| TAP2 R | ACTGCATCCTGGATCTCCC |
| STAT1 F | GAGCTTCACTCCCTTAGTTTTGA |
| STAT1 R | CACAACGGGCAGAGAGGT |
| STAT3 F | TCAAGACCTTCAGCTCCAAG |
| STAT3 R | TGACGCTGAGCGTGAAGAAG |
| CXCL10 F | GTGGCATTACAAGGAGTACCTC |
| CXCL10 R | TGATGGCCTTCGATTCTGGATT |
| B2M F | GGCATTCCTGAAGCTGACA |
| B2M R | CTTCAATGTCGGATGGATGAAAC |
| 18s F | GTAACCCGTTGAACCCCATT |
| 18s R | CCATCCAATCGGTAGTAGCG |
